# A threonine-to-aspartate ACA codon reassignment in uncultivated Acidimicrobiales bacteria

**DOI:** 10.64898/2026.05.03.722492

**Authors:** Fadel Fakih, Yekaterina Shulgina, Julius Lukeš, Anzhelika Butenko

## Abstract

The list of alternative genetic codes continues to grow, yet sense-to-sense codon reassignments remain very rare. Here we analyze a new sense-to-sense codon reassignment in bacteria, where the ACA codon, which canonically specifies threonine, has been predicted to encode aspartate. This so far unprecedented genetic code alteration is confined to a clade within the RAAP-2 family of the order Acidimicrobiales (phylum Actinomycetota). Validated by multiple alignments of highly conserved proteins, the reassignment is further supported by the presence of a tRNA_UGU_ that lacks the G1:C72 threonylation identity element, suggesting that this tRNA is charged with aspartate instead of threonine. Our findings not only expand the set of known natural departures from the canonical genetic code by a novel type of sense-to-sense reassignment but also implicate the ACA codon that was so far considered invariably conserved in its meaning.

## INTRODUCTION

The genetic code is a fundamental unifying feature of all extant life which is, as everything in the living domain, subject to tinkering (1). Despite being extremely ancient, conserved and refractory to changes, about 60 unique alterations of the genetic code have so far been documented in viruses, archaea, bacteria and the nuclear and organellar genomes of eukaryotes (2). While some codons are known to have only the canonical meaning, other codons are particularly prone to reassignment (2).

Probably due to their relative rarity and thus limited damage caused by the code switch, stop codons excel in this respect, sometimes facilitated by the presence of remaining dedicated termination codon and/or by context-dependent decoding (3). Moreover, stop codon reassignments are inherently easier to detect because they manifest as in-frame stop codons in predicted protein sequences when the respective coding sequences are conceptually translated using the canonical genetic code (4). Indeed, genetic code alterations involving sense codons are usually detected by dedicated programs, such as Codetta (5), and validated experimentally by comparing nucleotide sequences to proteomic mass spectrometry data. Initially, sense-to-sense reassignments were known only in mitochondria (6) and plastids (7), where they are accompanied by the loss of a dedicated transfer RNA (tRNA) (2). However, the view that this dramatic code alteration is restricted to organellar compartments was challenged by the discovery of nuclear sense-to-sense switches in yeast (8,9), which was followed by several distinct cases unearthed in various bacteria (5).

At the molecular level, codon reassignment typically involves alterations of certain components of the translation machinery. Both tRNAs and their respective aminoacyl-tRNA synthetases (aaRS) play a pivotal role in codon reassignment (10). While we still know surprisingly little about the specific relationships between particular amino acid substitutions in aaRSs and the corresponding alterations of the genetic code (1), much information is available regarding changes to tRNAs that alter their decoding capacity (11). These include mutations, nucleotide editing and modification within the anticodon, and the unpinning of the top base pair in the anticodon stem (2). Other variations outside of the anticodon arm can expand tRNA decoding through structural changes that increase flexibility of the tRNA body and promote non-canonical pairings at the wobble position (12). Sense codon reassignments, in particular, require new combinations of codons and amino acids. This can be achieved either by changing which codons a tRNA recognizes or by altering how a tRNA is aminoacylated. Codon-anticodon pairing may be modified through the mechanisms described above. As for the aminoacylation, aaRSs select the appropriate tRNAs by recognizing specific sequence and structural features known as identity elements (13). For many aaRSs, the tRNA anticodon is an important identity element, and as a result, reassignments that involve aaRSs that do not bind the anticodon are more common (10).

Sense-to-sense reassignments can be especially disruptive, as they replace one amino acid with another across all codon positions in the proteome (2,14). The magnitude of disruption depends on the frequency of the respective codon in the genome and biochemical nature of the two amino acids involved (15). Even single substitutions between chemically similar amino acids can impact the fitness of an organism, mainly through effects on protein folding, stability, and translation efficiency (16). Substitution between chemically distinct amino acids may lead to even more deleterious consequences, rendering them unlikely to be retained by evolution (17). This may contribute to why sense-to-sense reassignments are rarer than stop-to-sense ones. However, these proteome-wide costs can be greatly reduced under the codon capture model, where the codon is first driven to rarity or loss, thereby minimizing the immediate cost of reassignment before the codon later re-appears with a new meaning (18). It remains unclear why threonine codon reassignments, other than the one described here, have not been identified, given that GC content is a major determinant of threonine codon usage bias in bacteria as well as in archaea and eukaryotes (19), a pattern that is compatible with the codon capture model of the genetic code reassignment.

A switch from threonine to aspartate would represent a dramatic, non-conservative change. Threonine is a polar, uncharged amino acid with a hydroxyl side chain that frequently participates in hydrogen bonding and can be a site of phosphorylation (20). Aspartate, by contrast, is acidic and carries a negatively charged carboxylate group at physiological pH, commonly participating in salt bridges, metal coordination, and the active sites of enzymes (21,22). Replacing threonine with aspartate at multiple sites would thus alter the hydrogen-bonding networks (23). Replacement of the hydroxyl group-containing amino acids (threonine, serine, or tyrosine) with negatively charged residues (aspartate or glutamate) is commonly used to mimic site-specific protein phosphorylation (24). However, these phosphomimetic substitutions often only partially replicate authentic phosphorylation or even mimic the lack of it, as they do not fully reproduce the size or charge of a phosphate group (25,26). Thus, the large biochemical difference between threonine and aspartate would strongly constrain any reassignment route involving persistent ACA codons at ancestral threonine sites, although this constraint would be relaxed under a codon-capture scenario in which ACA had largely disappeared before reassignment.

Recently, we performed genetic code inference on 42,109 representative bacterial genomes from the Genome Taxonomy Database (GTDB) (27) and predicted a potentially novel reassignment within a clade of the bacterial family provisionally designated as the RAAP-2 of the order Acidimicrobiales (phylum Actinomycetota) (28). In parallel, this reassignment was identified using a new genetic code analysis software applied to a large dataset of prokaryotic genome assemblies (29). RAAP-2 clade brings together a group of deep-branching, uncultivated bacteria identified through genome-resolved metagenomics (30). Similar to other members of this order, RAAP-2 bacteria have streamlined genomes with elevated GC content and a genetic repertoire suggesting a high metabolic versatility, including an overrepresentation of transcriptional regulators and signal-transduction systems relative to their genome size (31). In this work, we analyze the genetic background of the threonine-to-aspartate ACA reassignment in a monophyletic lineage comprising 20 available genomes within the RAAP-2 clade. Using alignments of conserved orthologous proteins, structural analysis of tRNAs, and phylogenetics we provide multiple lines of evidence supporting this reassignment and explore its molecular and evolutionary basis.

## RESULTS

### A newly identified threonine-to-aspartate ACA reassignment in RAAP-2 bacteria

In the course of our previous analysis of tRNA repertoires and features underlying stop-to-tryptophan UGA reassignments in bacteria (28), we observed indications in the results of the Codetta software (5,32) used for genetic code inference that a novel threonine-to-aspartate ACA codon reassignment may occur in a subset of species within the RAAP-2 clade of the bacterial phylum Actinomycetota. This reassignment was recently independently identified using a different genetic code inference algorithm, providing further support for its validity (29). Here, we expanded the bacterial dataset by incorporating 260 representatives of the RAAP-2 clade (Suppl. table 1). Genetic code prediction by Codetta on these genomes clearly identified threonine-to-aspartate ACA reassignment in 20 bacterial species comprising the genera provisionally designated as JAJYVT01 (16 species), DAHUZJ01 (1 species), DASZUK01 (1 species) and JAJYWF01 (2 species) in the GTDB release 226 (Fig. 1A; Suppl. table 1). Across these species, the inference of ACA as aspartate by Codetta was based on 245 to 635 Pfam consensus columns (Suppl. table 1), which consistently positioned ACA codons at conserved aspartate sites within protein sequences. Two of the 20 genomes received a low confidence Codetta prediction (Fig. 1A). In both cases, however, the contig-level signal was largely consistent with threonine-to-aspartate ACA reassignment, with all informative contigs supporting the reassignment except for a single contig predicted to retain standard ACA decoding. Notably, unlike the contigs supporting the reassignment, this contig yielded top BLAST hits to Acidimicrobiales outside the RAAP-2 genomes analyzed here, suggesting that its presence within the RAAP-2 assembly likely reflects incorrect contig binning. Codetta uses Pfam alignments to identify conserved protein domains and infer the most likely translation for each codon, and this prediction may be subject to artifacts if there is a large discrepancy in the composition of the genome being analyzed and the Pfam models. Therefore, to confirm that the Codetta predictions were not affected by the high GC content of these genomes (median of 70.7%), we repeated the analysis using a custom version of Pfam built only from species with high GC content and found that the predicted threonine-to-aspartate ACA reassignment remained.

**Fig. 1.**
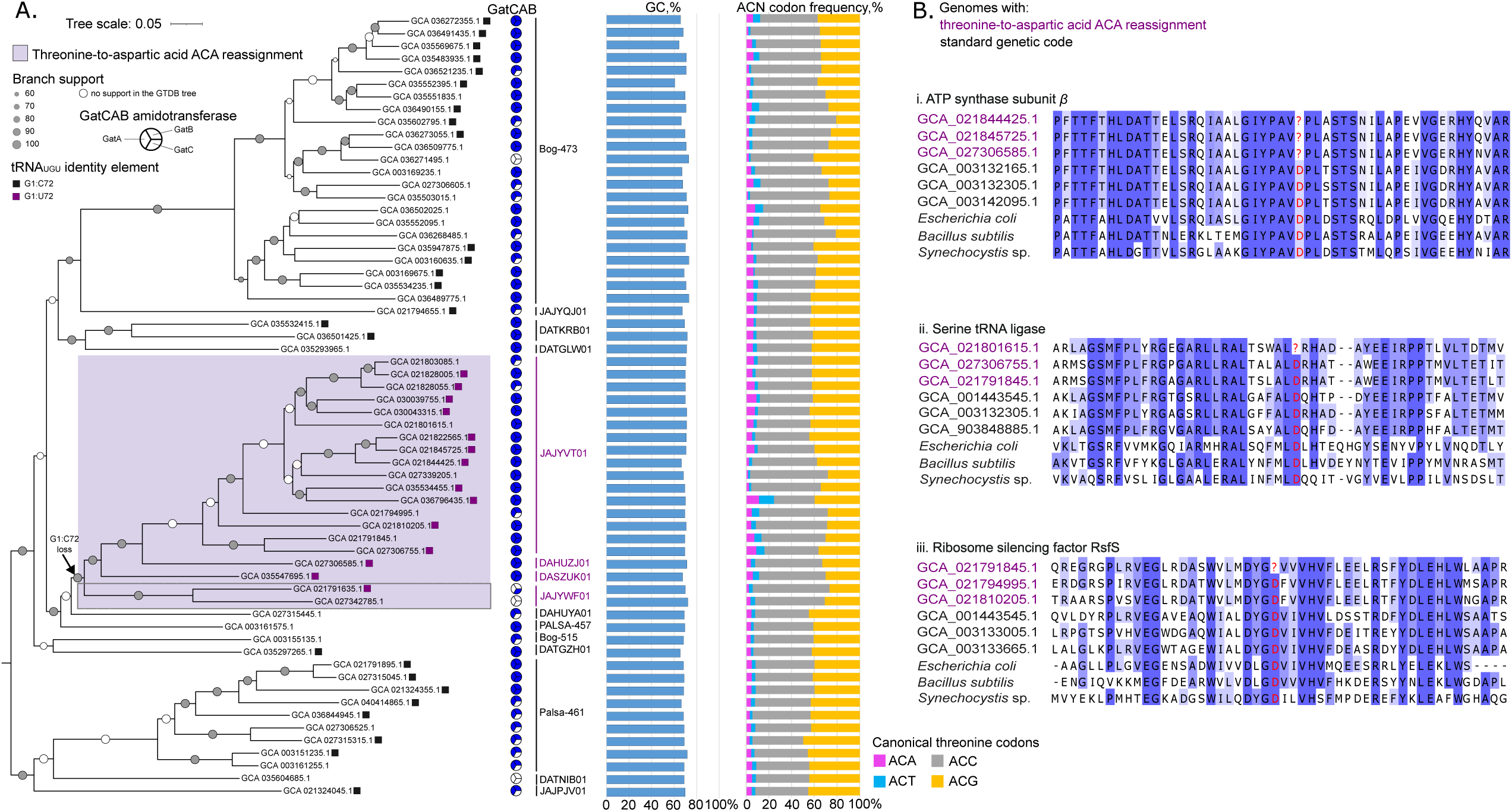
Threonine-to-aspartate ACA reassignment in the RAAP-2 clade. **(A**) Maximum-likelihood phylogenomic tree encompassing 20 representatives with the threonine-to-aspartate ACA reassignment (on the purple background) and 42 of their closest relatives with the standard genetic code. Two genomes with low-confidence Codetta predictions (see Results for details) are highlighted with a grey outline. Horizontal line indicates the number of substitutions per site. Branch support values represent the results of the Shimodaira-Hasegawa-like aLRT test; the support values for branches missing in the Genome Taxonomy Database (GTDB) are shown as white circles. The presence and absence of the threonine identity elements in tRNA_UGU_ is depicted with black and violet boxes, respectively. The absence of the box indicates that the tRNA_UGU_ gene was not identified. Pie charts indicate the presence (blue)/ absence (white) of the genes encoding the subunits GatA, GatB, and GatC of the GatCAB amidotransferase, performing the synthesis of tRNA^Asn^. Bar plots show the GC percentage and the proportion of the threonine codons (ACA, ACT, ACC, and ACG) in the RAAP-2 genomes. Genus names are indicated alongside the vertical lines, with those corresponding to bacteria exhibiting the genetic code reassignment shown in violet. (**B**) Alignments of the conserved domains for ATP synthase subunit β, serine tRNA ligase, and ribosomal silencing factor RsfS. Residues are shaded according to percentage identity, with darker shading indicating higher conservation.

To further validate the Codetta prediction, we identified homologs of proteins from bacteria with the ACA reassignment in the dataset of benchmarking universal single-copy orthologs (BUSCOs) (33). We aligned 20 proteins containing an ACA codon within their coding sequence in at least one species in the putatively reassigned clade to 1,000 homologues from phylogenetically diverse bacteria. In three cases, the ACA codon aligned with a conserved aspartate present in ≥70% of the reference sequences; in the remaining 17 cases, the corresponding position was still most frequently occupied by aspartate, albeit at a frequency below 70%. For example, an ACA codon aligned with a conserved aspartate immediately downstream of the catalytic nucleotide-binding domain of the ATP synthase subunit β (Fig. 1B) (34). This is a position occupied by aspartate in 97% of homologs in bacteria with the standard genetic code. In the second example, an ACA site in the gene encoding serine tRNA ligase (SerRS) occurs at a conserved position (aspartate in 81.2% of the reference sequences) upstream to motif I within the catalytic domain (Fig. 1B) (35,36). A third example comprises the ribosome silencing factor (RsfS), which in a reassigned genome bears the ACA codon at a conserved aspartate position (present in 73.1% of the reference homologues) within the DNA polymerase *β* nucleotidyltransferase superfamily protein (37). Notably, in some of the reassigned genomes, the conserved aspartate residue in SerRS and RsfS sequences is decoded by a canonical aspartate codon, rather than by ACA, suggesting that these may be synonymous aspartate codons in this clade.

To provide a robust evolutionary framework, we built a phylogenomic tree using 492 conserved proteins encoded by single-copy genes in 62 genomes in the RAAP-2 clade, incorporating species with the reassigned genetic code and their closest relatives (according to the phylogenomic tree available from GTDB) with the standard genetic code (Suppl. table 2). The concatenated alignment produced a well-resolved maximum-likelihood tree (Fig. 1A). Notably, all genomes with this predicted reassignment form a strongly supported clade nested within the RAAP-2 lineage in both our phylogenetic analysis and the GTDB tree (Fig. 1A). This indicates that the threonine-to-aspartate ACA reassignment had occurred in the common ancestor of the genera JAJYVT01, DAHUZJ01, DASZUK01, and JAJYWF01.

Next, we analyzed codon usage across the RAAP-2 clade (Fig. 1A; Suppl. table 2). Species with the threonine-to-aspartate ACA reassignment, together with their closest relatives with the standard genetic code (Fig. 1A), display relatively high GC content (median 69.8%). Such elevated GC content is associated with significantly lower usage of the AT-rich threonine codons (ACT and ACA), compared to the GC-rich codons (ACC and ACG) (two-sided Mann–Whitney U test, *p*-value < 0.0001) (Fig. 1A). The GC-driven rarity of ACA in both the taxa with an ACA reassignment and their closest relatives with the standard genetic code likely facilitated the emergence of the reassignment in accordance with the codon capture scenario (18). Notably, ACA is used significantly more frequently in the genomes demonstrating the reassignment (median of 6.3% of all threonine codons) than in the standard code relatives (4.3%) (*p* < 0.0001; Suppl. table 2), suggesting a trend toward increasing ACA usage following the reassignment.

### tRNA_UGU_ identity features are consistent with a mechanistic explanation for the reassignment

In search for a mechanistic explanation of the discovered reassignment, we focused on the tRNAs and aaRSs in genomes with the reassigned code and their closest relatives with the standard genetic code (Fig. 1A). We identified a tRNA_UGU_ gene, which is anticipated to read ACA codons, in 14 out of 20 genomes with the threonine-to-aspartate ACA reassignment, while genomes where it was absent have genome completeness estimates ranging from 58.5 to 91.1% (Suppl. table 2). Notably, the identified tRNA_UGU_s lack an important tRNA^Thr^ identity element, G1:C72 (38), and instead have a G1:U72 (Fig. 2A), in contrast to the tRNA^Thr^_UGU_ from the RAAP-2 bacteria with the standard genetic code (Fig. 2B). The G1:C72 identity element is also conserved in tRNA^Thr^_CGU_ (Fig. 2C) and tRNA^Thr^_GGU_ sequences (Fig. 2D) within all representatives of the RAAP-2 clade, regardless of the genetic code. At a broader scale, C72 appears to be highly conserved among bacterial tRNA^Thr^: in a dataset of 110,692 bacterial tRNA^Thr^ genes (28), C72 is present in 110,206 sequences (99.56%), whereas U72 is extremely rare, being present in only 75 sequences (∼0.07%) scattered across multiple bacterial phyla, underscoring how exceptional is the G1:U72 pair observed in the reassigned clade.

**Fig. 2.**
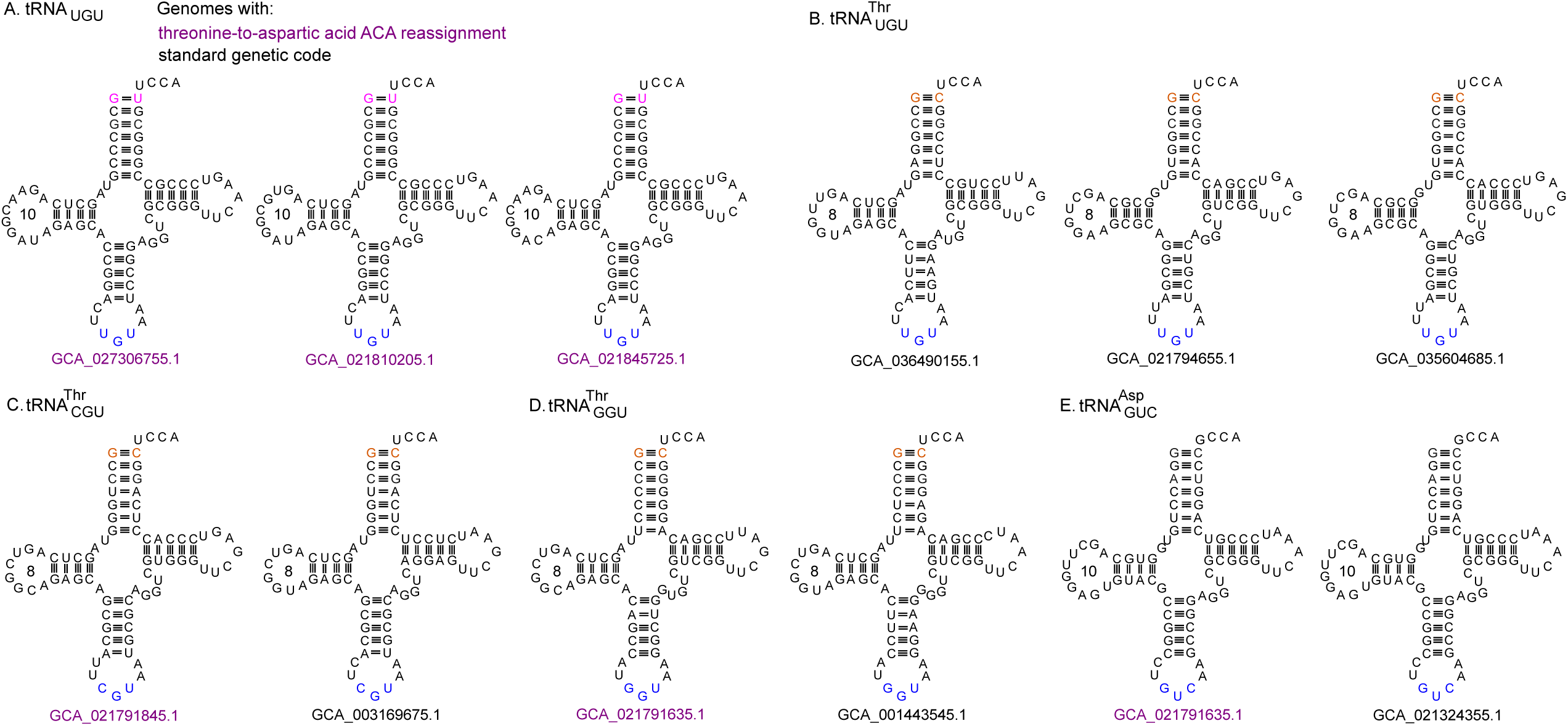
tRNA_UGU_ features are consistent with the threonine-to-aspartate identity switch. (**A**) Secondary structure of tRNA_UGU_ from reassigned genomes carrying G1:U72 base pair (magenta) at the top of the acceptor stem (loss of C1:G72 threonylation identity). In contrast, tRNA^Thr^_UGU_ (**B**), tRNA^Thr^_CGU_ (**C**), and tRNA^Thr^_GGU_ (**D**) from genomes with the standard genetic code retain the canonical G1:C72 pair (orange). tRNA_UGU_ from the reassigned genomes show structural similarities (*e.g*., 10-nt D-loop) to tRNA^Asp^_GUC_ (**E**). Anticodon nucleotides are shown in blue. The genome accession number corresponding to each tRNA sequence is indicated below its secondary structure.

Importantly, loss of the tRNA^Thr^_UGU_ G1:C72 identity element occurred in the common ancestor of the predicted threonine-to-aspartate ACA reassignment (Fig. 1A). The secondary structure-based alignment of tRNA_UGU_, tRNA^Thr^_CGU_, tRNA^Thr^_GGU_, and tRNA^Asp^_GUC_ from the reassigned genomes and those from RAAP-2 species that use the standard genetic code (Suppl. Fig. 1A) revealed that tRNA_UGU_ in the reassigned genomes is on average slightly more similar to tRNA^Asp^_GUC_ than to other tRNA^Thr^ isoacceptors (65.44 ± 1.39% vs 62.43 ± 2.56% pairwise identity, respectively; *p*-value < 0.01). In contrast, the corresponding tRNA^Thr^_UGU_ from the outgroup with the standard genetic code are more similar to other tRNA^Thr^ isoacceptors than to tRNA^Asp^_GUC_ (68.42 ± 4.77% vs 51.18 ± 2.16%, respectively; *p*-value < 0.01). The major determinants of tRNA^Asp^_GUC_ identity, namely the anticodon bases G34, U35, C36, and the discriminator base G73, are absent in tRNA_UGU_, while they are present in tRNA^Asp^_GUC_ in both bacteria with an ACA reassignment and those with the standard genetic code. Another aspartate tRNA identity element, the base G10 in the D-stem (39) is present in tRNA_UGU_ and tRNA^Asp^_GUC_ in both bacteria with an ACA reassignment and those with the standard genetic code, while also being nearly universally conserved in 43,485 (99.26%) bacterial tRNA^Thr^_UGU_ sequences in the dataset from Fakih *et al*. (28). Thus, it is unlikely to contribute to the ACA reassignment reported here. Notably, tRNA_UGU_s from bacteria with an ACA reassignment share a conserved 10-nucleotide (nt) long D-loop (Fig. 2A), compared to 7- or 8-nt D-loops in other tRNA^Thr^ (Fig. 2B-D). A similar 10-nt D-loop is also present in tRNA^Asp^_GUC_ of both reassigned and not reassigned genomes (Fig. 2E). In a tRNA phylogenetic tree constructed from tRNA^Thr^ and tRNA^Asp^ sequences from RAAP-2 genomes and other Acidomicrobiales species, the tRNA_UGU_ sequences from the reassigned clade group separately from either tRNA^Thr^ or tRNA^Asp^ sequences (Suppl. fig. 1B), precluding any clear conclusions about shared evolutionary origin or aminoacylation identity.

To further investigate the origin of tRNA_UGU_ in the reassigned clade, we performed a synteny analysis incorporating 9 genomes with an ACA reassignment and their closest relatives with the standard genetic code. While syntenic within the reassigned genomes (Suppl. fig. 2). tRNA^Thr^_UGU_s occur within a different genomic context in the RAAP-2 genomes with the standard genetic code, preventing any conclusion about the origin of tRNA_UGU_ in the reassigned genomes (Suppl. fig. 2). Genomic regions in bacteria with the standard genetic code lack tRNA genes at the locus syntenic to the tRNA_UGU_ locus in bacteria with an ACA reassignment, hindering synteny-based inference of the evolutionary history of the tRNA genes of interest in RAAP-2.

We wondered whether an aaRS gene duplication may be behind an aminoacylation system supporting the ACA reassignment. However, we did not identify any aaRS genes resulting from gene duplication events and encoding a dedicated AspRS potentially capable of aminoacylating tRNA_UGU_ in bacteria exhibiting the ACA reassignment. Thus, the most likely candidate for aminoacylating such tRNA_UGU_ is AspRS with an expanded tRNA specificity. The absence of asparaginyl-tRNA synthetase (AsnRS), together with the presence of the GatCAB amidotransferase complex (Fig. 1A) in the analyzed RAAP-2 bacteria indicate that they use an indirect tRNA-dependent pathway (40) for Asn-tRNA^Asn^ synthesis. In this pathway, a non-discriminating (ND) AspRS first generates Asp-tRNA^Asn^, which is subsequently converted to Asn-tRNA^Asn^ by GatCAB (41). Accordingly, in these bacteria, AspRS is expected to be non-discriminating, recognizing both tRNA^Asp^ and tRNA^Asn^ (40), and, thus, inherently exhibiting expanded tRNA specificity.

In addition, mutations at certain amino acid positions (*e.g*., within the anticodon-binding domain) of the ND AspRS have been demonstrated to modulate its tRNA specificity (40,42). The alignment of ND AspRSs from the RAAP-2 bacteria with the ACA reassignment, their closest relatives with the standard code, as well as from *Thermus thermophilus* (which has a well-studied ND AspRS) (40) revealed some mutations within the OB-fold anticodon-binding region (Suppl. fig. 3). The core anticodon readout residues Phe33, Gln44, and Glu76 remain conserved, whereas the entire RAAP-2 lineage (both bacteria with an ACA reassignment and those with the standard genetic code) carries a P72T substitution at the L1-loop implicated in U36 tolerance, which indicates a potentially lower tRNA specificity of AspRS in the analyzed RAAP-2 bacteria (40). In addition, there are clear differences in the conserved positions within the AspRS anticodon-binding domain between the reassigned RAAP-2 bacteria and those with the standard genetic code (Suppl. fig. 3). Strikingly, there is a V66E substitution in bacteria with an ACA reassignment relative to both *T. thermophilus* and RAAP-2 bacteria with the standard genetic code (Suppl. fig. 3), as well as M65T substitution (this position in *T. thermopilus* corresponds to a gap) in the S4 region of the anticodon-binding domain (40). Combined, these results suggest that both tRNA_UGU_ and AspRS in the reassigned clade have acquired features potentially compatible with AspRS-mediated charging of tRNA_UGU_, providing a plausible mechanistic basis for the ACA reassignment.

## DISCUSSION

To the best of our knowledge, only 19 out of 64 codons of the canonical genetic code have been found recoded in at least a single species across the three domains of life (2). Here we analyze ACA, the 20^th^ reassigned codon in this slowly growing collection. In bacteria, only seven distinct instances of the genetic code reassignment have been documented so far (2). Notably, UGA is the only stop codon subjected to the stop-to-sense reassignments, with the switch to tryptophan evolving multiple times independently, such as in Mycoplasmatales and *Candidatus* Zinderia spp. (43,44). The other amino acid specified by UGA in Gracilibacteria and Absconditabacteria is glycine (5). On the other hand, all the sense-to-sense reassignments affect arginine codons. Both CGR codons specify tryptophan in Absconditabacteria, while in *Anaerococcus* and *Candidatus* Onthovivens, the same reassignment is confined only to the CGG codon (2). Two additional alternative codes are represented by CGG and AGG specifying glutamine and methionine in *Peptacetobacter* and *Enterosoma*, respectively (5). The threonine-to-aspartate ACA reassignment represents only the second known instance of codon reassignment in Actinomycetota, in addition to the recently described stop-to-tryptophan UGA reassignment that has occurred at least twice independently within the family Eggerthellaceae (45).

A monophyletic lineage within the RAAP-2 clade of the order Acidimicrobiales exhibits a so far unprecedented reassignment of the ACA codon from threonine to aspartate (28,29). In the absence of mass spectrometry data from these uncultivated bacteria, this switch is supported by several lines of indirect evidence, namely alignments with the conserved homologues from bacteria with the standard genetic code, tRNA identity signatures, a likely expanded ND AspRS specificity, and an apparent monophyly.

Multiple alignments of conserved single-copy orthologues strongly support ACA being decoded as aspartate in this subset of RAAP-2 genomes, including in proteins where ACA occurs within functionally and structurally important domains. The first such case is represented by the ACA codon located at a conserved Asp residue close to motif I within the catalytic domain of SerRS, which is required for ATP binding and catalysis (35,36). Another functionally constrained case comprises RsfS, where in the reassigned genomes the ACA codon occurs at a conserved aspartate residue within the DNA polymerase β nucleotidyltransferase superfamily protein. This domain includes a conserved catalytic core built around a Aspx[Asp/Glu] motif and a third carboxylate residue (37). Despite the broad functional and sequence diversity across this protein family, the Aspx[Asp/Glu] motif remains essential for coordinating two divalent metal ions critical for nucleotide transfer and is thus highly conserved (37). In the third case, the ACA codon corresponds to a highly conserved aspartate residue in ATP synthase subunit β, where it plays a key role in interactions with other subunits of the ATP synthase complex (46).

Features of the tRNA_UGU_ offers a plausible mechanistic explanation for the reassignment. Mutation of the C1:G72 base pair of tRNA^Thr^ in of *E. coli* abolishes threonylation (38). Indeed, among the RAAP-2 bacteria, only the reassigned clade tRNA_UGU_ lacks a major threonylation identity element, the G1:C72 pair, and possesses the G1:U72 instead. Importantly, the G1:C72 base pair is virtually universally conserved in tRNA^Thr^ bacteria, which may indicate selection pressure on this functionally significant position. Moreover, in the reassigned RAAP-2 bacteria, tRNA_UGU_ is slightly more similar to tRNA^Asp^_GUC_ than to other tRNA^Thr^ species, consistent with several possible evolutionary scenarios, including an identity switch from tRNA^Thr^_UGU_ to tRNA^Asp^_UGU_, duplication of a tRNA^Asp^ gene followed by an anticodon mutation in one of the copies, or the evolution of another tRNA species that acquired both ACA codon recognition and the features enabling it to be charged with aspartate. Yet, because the tRNA_UGU_ genes in the reassigned genomes are not syntenic with the corresponding tRNA loci in the outgroup, their evolutionary origin cannot be determined. The reassignment may have been facilitated by the relaxed specificity of the ND AspRS. Rather than evolving a dedicated enzyme, the RAAP-2 bacteria appear to use a ND AspRS with an expanded tRNA specificity, with substitutions in the anticodon-binding domain potentially enabling the aspartinylation of the putative tRNA^Asp^_UGU_. Together, the presence of mutations at the amino acid positions critical for the interaction with the anticodon in the AspRS, as well as deviations to the tRNA_UGU_ identity elements, suggest that the reassignment is enabled by the tRNA identity switch.

Our phylogenomic analyses show that the threonine-to-aspartate ACA reassignment in RAAP-2 is monophyletic, hence has evolved in the analyzed bacteria only once, and this unique event apparently did not prevent diversification of the affected lineage. In contrast to other codon reassignments in bacteria that evolved multiple times independently, such as the stop-to-tryptophan UGA switch (2), this event is so far known to have occurred only once in bacteria (28,29). The codon usage analysis shows that ACA was driven to rarity already in the ancestor of the reassigned bacteria, as judged by the high GC content in the genomes of their closest relatives. The available data is consistent with a codon capture scenario, in which the depletion of ACA due to increasing GC bias prior to the reassignment was followed by re-expansion of this codon following the reassignment.

A notable feature of the reassignment described here is that it involves a single NNA codon (ACA). In most documented cases of sense codon reassignment, either NNG codons are affected, owing to their decoding by dedicated C34 tRNAs that can be altered without impacting other synonymous codons (47), or the reassignment results in the merger of codon boxes, as exemplified by the well-known stop-to-tryptophan UGA reassignment that expands tryptophan decoding to UGR codons (48). In contrast, isolated reassignment of an NNA codon is rare, likely because U34-containing tRNAs can decode NNG codons *via* wobble pairing, in addition to NNA codons. In the case reported here, tRNA_UGU_ in bacteria with threonine-to-aspartate ACA reassignment, has the potential to mistranslate threonine ACG codons as aspartate *via* wobble interaction. Considering a quite high proportion of ACG codons (median of 41.6%) in bacteria with the ACA reassignment, such mistranslation could have a dramatic effect, unless tRNA_UGU_ is expressed at low levels and the resulting mistranslation events occur only infrequently. Hence, there should be a mechanism to prevent ACG misreading as aspartate by tRNA_UGU_ in the reassigned genomes. Achieving differential decoding of NNA and NNG therefore requires additional constraints, most plausibly at the level of wobble base modification. In bacteria, U34 is frequently modified (*e.g*., 5-methylaminomethyl-2-thiouridine and carboxymethylaminomethyl-2-thiouridine), which promotes preferential U-A pairing while restricting the U–G wobble interactions, thereby biasing decoding toward NNA codons (47,49). Such modifications could, in principle, enable selective reassignment of ACA while minimizing mistranslation of ACG.

In summary, we provide multiple lines of evidence to support a threonine-to-aspartate ACA reassignment in Acidimicrobiales. This is a novel sense-to-sense codon reassignment, expanding the still rather narrow collection of this type of genetic code alterations. Our study highlights the role of plasticity in defining tRNA identity in driving codon reassignments. Finally, our findings are informative for the burgeoning generation of the artificial genetic codes by documenting that the wholesale threonine-to-aspartate ACA switch is, at least in some bacteria, compatible with survival.

## MATERIALS AND METHODS

### Phylogenomic analysis

Phylogenetic placement of 20 bacteria exhibiting the threonine-to-aspartate ACA reassignment was initially evaluated using a phylogenomic tree available in the GTDB release 226. We then selected an additional set of 42 genomes (Suppl. table 2) according to the GTDB tree representing a well-supported clade (bootstrap support of 95%) within RAAP-2 and identified them as the closest known relatives of the reassigned bacteria. To verify monophyly of the lineage demonstrating an ACA reassignment, we inferred orthologous groups of proteins with OrthoFinder v.2.5.5 (options: -S blast, -T iqtree, -M msa, and all other settings left at default values) (50) using a dataset composed of RAAP-2 genomes with the genetic code reassignment and their nearest relatives (Suppl. table 2). Phylogenomic analysis was conducted on 492 proteins encoded by single-copy genes present in at least 68% of the analyzed genomes. The phylogenomic tree was inferred using IQ-TREE v.3.0.1 (51) with automatic model selection, and branch support values were calculated using the Shimodaira-Hasegawa-like aLRT test. Prior to the tree reconstruction, protein sequences were aligned with MAFFT v.7.505 employing the L-INS-i algorithm (52) and trimmed using trimAl v.1.4.rev15 (53) with the -gt 0.8 option. The tree was visualized with Interactive Tree Of Life (iTOL) web server (54).

### Genetic code inference and codon usage analysis

Genetic code predictions of 260 genomes of the RAAP-2 clade (Suppl. table 2) were performed using Codetta v.2.0 with default settings (32). A custom high GC version of Pfam 35.0 (55) was constructed by subsetting the Pfam full alignments to sequences originating from species with a genomic GC content > 0.65, and building profile HMMs for all models using the HMMER v.3.3.2 program hmmbuild (56). Codetta analyses were repeated as before, specifying the custom Pfam database with the -p option. The subset of 62 RAAP-2 genomes (Suppl. table 2) was annotated using Prokka v.1.14.6 (57). Codon count and GC content were computed using EMBOSS cusp v.6.6.0.0 (58). The statistical significance of differences in GC content and codon usage between the genomes with threonine-to-aspartate ACA reassignment and those with the standard genetic code was tested using a Mann-Whitney U test in GraphPad Prism v.10.6.1 (59).

### tRNA prediction, identity elements, and structural comparison

tRNA genes for the genomes listed in the Suppl. table 2 were predicted using ARAGORN v.1.2.41. tRNA^Thr^ isoacceptors and tRNA^Asp^_GUC_ were initially aligned using LocARNA v.1.9.1 (60) and manually refined. Sequence identity values were obtained with Clustal Omega v.1.2.4. and used to compare similarity patterns between the various tRNAs. The tRNA phylogenetic tree was constructed from tRNA^Thr^ and tRNA^Asp^ genes identified from all RAAP-2 genomes and ten additional Acidomicrobiales genomes, dereplicated to remove identical sequences and processed to remove CCA tails and mask the anticodon. The tree was constructed with MrBayes 3.2.7a (61) run for 50 million generations with sampling every 500 generations and default burn-in. Alignment columns were partitioned into stem and loop regions, modelled by separately parameterized GTR substitution models with gamma-distributed rate variation across sites and a proportion of invariable sites. The 3’ halves of stems were omitted from the phylogenetic analysis to avoid correlated information present in base paired regions. Synteny analysis was performed using Easyfig v.2.2.5 with default parameters (62).

### Homology searches and sequence alignments

The homologs of aaRSs and the GatCAB amidotransferase complex subunits were searched for by BLAST (BLASTp and tBLASTn) (63) with an *e*-value threshold of 1e-3. The homologs from *E. coli*, *Thermus thermophilus*, and *Helicobacter pylori* (Suppl. table 3) were used as queries, and the genome-derived proteomes and genome sequences of the 62 RAAP-2 bacteria listed in the Suppl. table 2 as the BLAST database. BUSCO searches were performed with BUSCO v.6.0.0 implemented in Galaxy platform with the OrthoDB v.12 (64). The homologs of proteins encoded by the genes with in-frame ACA codons in the RAAP-2 genomes with an ACA reassignment and those with the standard genetic code were extracted from BUSCO dataset and aligned with query sequences using MAFFT v.7.505 employing the L-INS-i algorithm. The AspRS homologs were aligned using the same method.

## SUPPLEMENTARY FIGURE LEGENDS

**Suppl. fig. 1.**
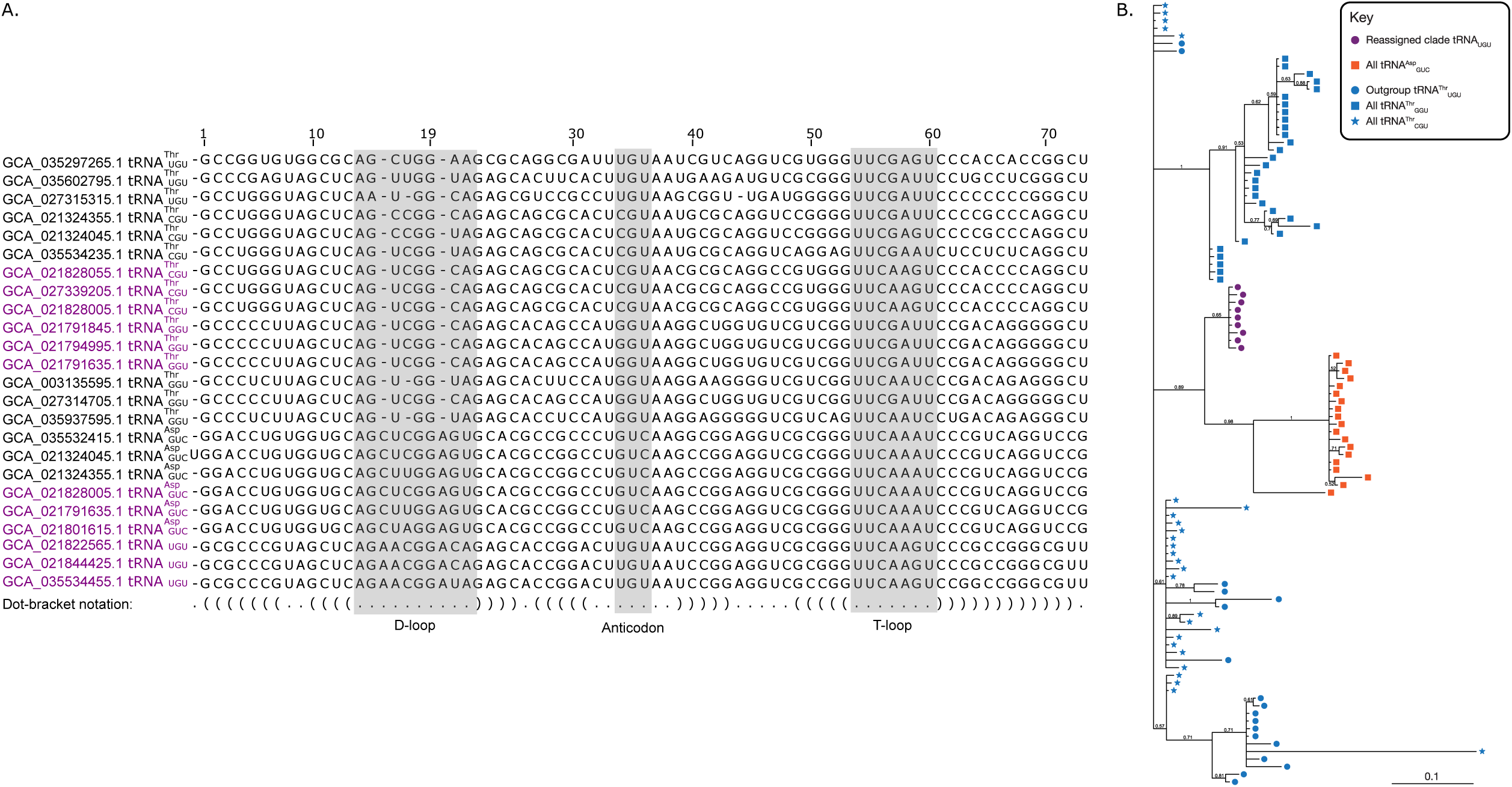
tRNA alignments. (**A)** Alignment of tRNA_UGU_, tRNA^Thr^, tRNA^Thr^, and tRNA^Asp^_GUC_ from reassigned and standard genetic code genomes. The numbering follows the Sprinzl system (65). Genome accessions for bacteria demonstrating the threonine-to-aspartate ACA reassignment are in violet. (**B)** Unrooted phylogenetic tree of tRNA_UGU_, tRNA^Thr^ and tRNA^Asp^ sequences from RAAP-2 and other Acidomicrobiales genomes. Branch values indicate the posterior probability of the branching. The branch length scale bar is in expected substitutions per site.

**Suppl. fig. 2.**
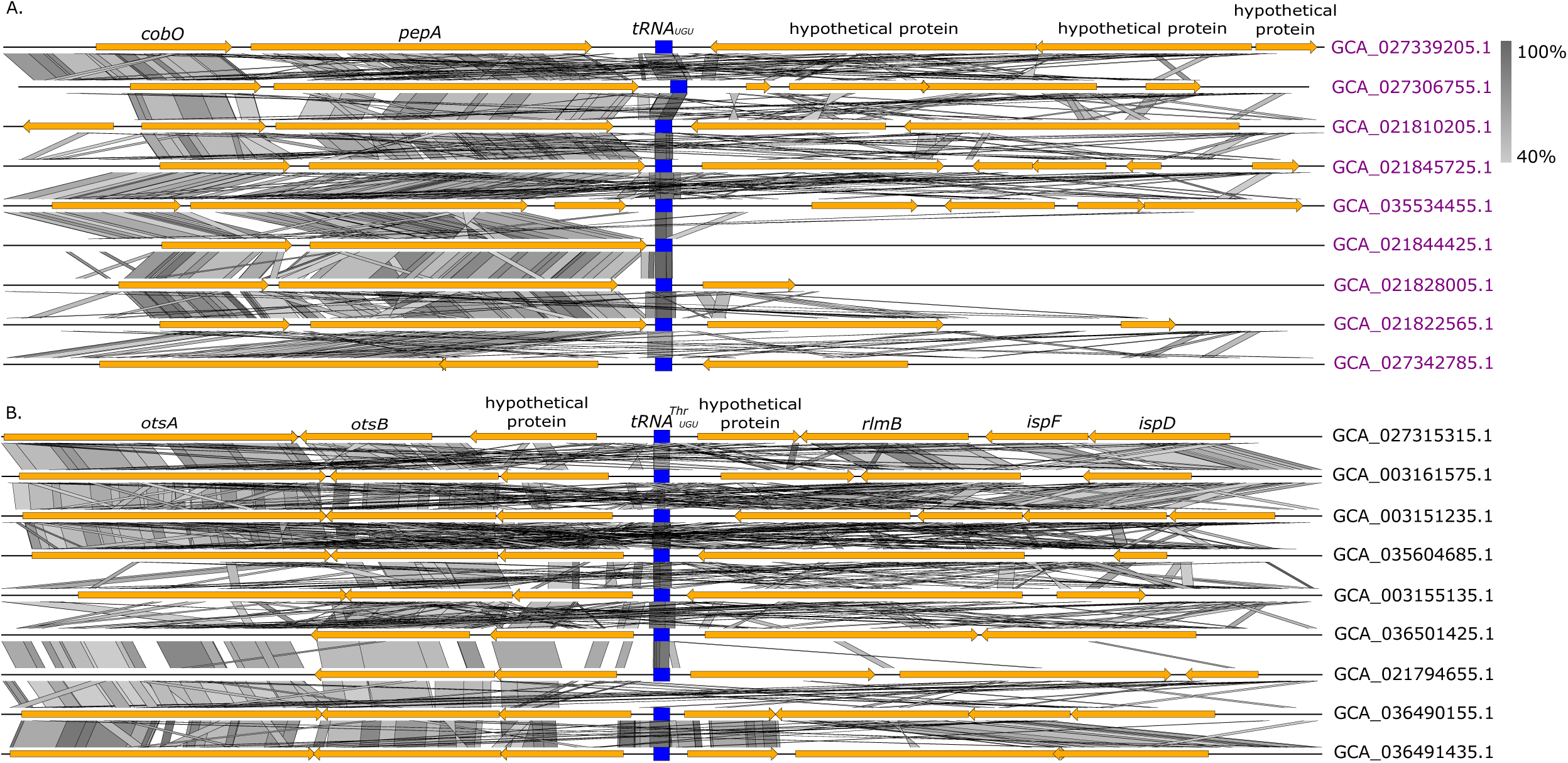
Synteny analysis showing the genomic context of tRNA_UGU_ genes in the reassigned genomes. (**A**) and their closest relatives with the standard genetic code. (**B**) (3,000 nucleotides to the right and left of the tRNA locus are shown). tRNA and protein-coding genes are shown as blue rectangles and orange arrows, respectively. Grey rectangles indicate similar genomic regions with the shade of grey reflecting the degree of similarity. Gene names: *cobO* - corrinoid adenosyltransferase; *ispD* - 2-C-methyl-D-erythritol 4-phosphate cytidylyltransferase; *ispF* - 2-C-methyl-D-erythritol 2,4-cyclodiphosphate synthase; *otsA -* trehalose-6-phosphate synthase; *otsB* - trehalose-6-phosphate phosphatase; *pepA* - aminopeptidase A; *rlmB* - 23S rRNA (guanosine^2251^-2’-O)-methyltransferase.

**Suppl. fig. 3.**
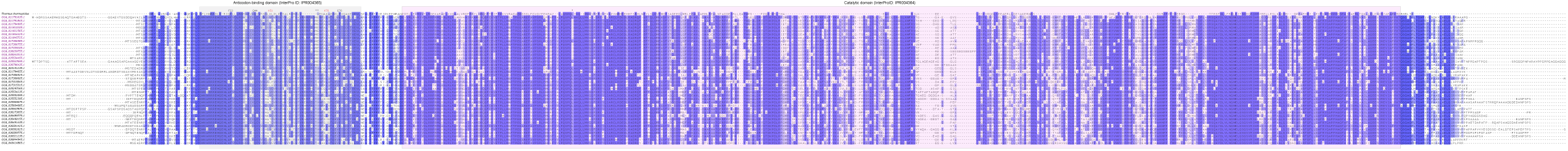
Sequence alignment of non-discriminating aspartyl-tRNA synthetases (ND AspRS) from the RAAP-2 bacteria with the threonine-to-aspartate ACA reassignment and those with the standard genetic code. AspRS of *Thermus thermophilus* is used as a reference. Anticodon-binding and catalytic domain are shown on grey and pink background, respectively. Genome accessions for bacteria with the threonine-to-aspartate ACA reassignment are in violet. ** - amino acid position with potential impact on the tRNA specificity that differs between the reassigned RAAP-2 bacteria and those with the standard genetic code.

## SUPPLEMENTARY TABLE LEGENDS

**Suppl. table 1**. Predicted genetic codes of the RAAP-2 clade bacteria.

**Suppl. table 2**. Results of the codon usage analysis in 20 RAAP-2 bacteria with the threonine-to-aspartate ACA reassignment (in red) and 42 of their closest relatives with the standard genetic code.

**Suppl. table 3**. Accession numbers for query sequences used for the identification of aaRS and GatCAB amidotransferase homologs in the RAAP-2 bacteria.

## REFERENCES

1. Mukai T, Lajoie MJ, Englert M, Söll D. Rewriting the genetic code. Annu Rev Microbiol. 2017 Sep 8;71:557–77. doi:10.1146/annurev-micro-090816-093247 PubMed PMID: 28697669; PubMed Central PMCID: PMC5772603.

2. Lukeš J, Eliáš M, Kachale A, van der Gulik PTS, Speijer D. Natural and artificial variations of the standard genetic code. Curr Biol. 2025 Nov 17;35(22):R1104–26. doi:10.1016/j.cub.2025.09.071.

3. Knight RD, Freeland SJ, Landweber LF. Rewiring the keyboard: evolvability of the genetic code. Nat Rev Genet. 2001 Jan;2(1):49–58. doi:10.1038/35047500.

4. Keeling PJ. Genomics: Evolution of the genetic code. Curr Biol. 2016 Sep 26;26(18):R851–3. doi:10.1016/j.cub.2016.08.005.

5. Shulgina Y, Eddy SR. A computational screen for alternative genetic codes in over 250,000 genomes. eLife. 2021 Nov 9;10:e71402. doi:10.7554/eLife.71402.

6. Sengupta S, Yang X, Higgs PG. The mechanisms of codon reassignments in mitochondrial genetic codes. J Mol Evol. 2007 Jun;64(6):662–88. doi:10.1007/s00239-006-0284-7 PubMed PMID: 17541678; PubMed Central PMCID: PMC1894752.

7. Matsumoto T, Ishikawa SA, Hashimoto T, Inagaki Y. A deviant genetic code in the green alga-derived plastid in the dinoflagellate *Lepidodinium chlorophorum*. Mol Phylogenet Evol. 2011 Jul;60(1):68–72. doi:10.1016/j.ympev.2011.04.010 PubMed PMID: 21530665.

8. Kawaguchi Y, Honda H, Taniguchi-Morimura J, Iwasaki S. The codon CUG is read as serine in an asporogenic yeast *Candida cylindracea*. Nature. 1989 Sep;341(6238):164–6. doi:10.1038/341164a0.

9. Yokogawa T, Suzuki T, Ueda T, Mori M, Ohama T, Kuchino Y, et al. Serine tRNA complementary to the nonuniversal serine codon CUG in *Candida cylindracea*: evolutionary implications. Proc Natl Acad Sci USA. 1992 Aug 15;89(16):7408–11. doi:10.1073/pnas.89.16.7408.

10. Kollmar M, Mühlhausen S. How tRNAs dictate nuclear codon reassignments: Only a few can capture non-cognate codons. RNA Biol. 2017 Jan 17;14(3):293–9. doi:10.1080/15476286.2017.1279785 PubMed PMID: 28095181; PubMed Central PMCID: PMC5367256.

11. Santos MAS, Moura G, Massey SE, Tuite MF. Driving change: the evolution of alternative genetic codes. Trends Genet. 2004 Feb;20(2):95–102. doi:10.1016/j.tig.2003.12.009 PubMed PMID: 14746991.

12. Schmeing TM, Voorhees RM, Kelley AC, Ramakrishnan V. How mutations in tRNA distant from the anticodon affect the fidelity of decoding. Nat Struct Mol Biol. 2011 Apr;18(4):432–6. doi:10.1038/nsmb.2003 PubMed PMID: 21378964; PubMed Central PMCID: PMC3072312.

13. Giegé R, Eriani G. The tRNA identity landscape for aminoacylation and beyond. Nucleic Acids Res. 2023 Feb 28;51(4):1528–70. doi:10.1093/nar/gkad007 PubMed PMID: 36744444; PubMed Central PMCID: PMC9976931.

14. Gomes AC, Miranda I, Silva RM, Moura GR, Thomas B, Akoulitchev A, et al. A genetic code alteration generates a proteome of high diversity in the human pathogen *Candida albicans*. Genome Biol. 2007 Oct 4;8(10):R206. doi:10.1186/gb-2007-8-10-r206.

15. Haig D, Hurst LD. A quantitative measure of error minimization in the genetic code. J Mol Evol. 1991 Nov;33(5):412–7. doi:10.1007/BF02103132 PubMed PMID: 1960738.

16. Parmley JL, Hurst LD. How do synonymous mutations affect fitness? Bioessays. 2007 Jun;29(6):515–9. doi:10.1002/bies.20592 PubMed PMID: 17508390.

17. Sun M, Stoltzfus A, McCandlish DM. A fitness distribution law for amino-acid replacements. bioRxiv [Preprint]. 2024 Oct 15;2024.10.11.617952. doi:10.1101/2024.10.11.617952 PubMed PMID: 39464166; PubMed Central PMCID: PMC11507765.

18. Yamashita T, Narikiyo O. Codon capture and ambiguous intermediate scenarios of genetic code evolution. arXiv [Preprint]. 2011 [cited 2025 Jul 12]. Available from: http://arxiv.org/abs/1110.5123. doi:10.48550/arXiv.1110.5123.

19. Novoa EM, Jungreis I, Jaillon O, Kellis M. Elucidation of codon usage signatures across the domains of life. Mol Biol Evol. 2019 Oct 1;36(10):2328–39. doi:10.1093/molbev/msz124.

20. Eswar N, Ramakrishnan C. Deterministic features of side-chain main-chain hydrogen bonds in globular protein structures. Protein Eng. 2000 Apr;13(4):227–38. doi:10.1093/protein/13.4.227 PubMed PMID: 10810153.

21. Craik CS, Roczniak S, Largman C, Rutter WJ. The catalytic role of the active site aspartic acid in serine proteases. Science. 1987 Aug 21;237(4817):909–13. doi:10.1126/science.3303334.

22. Dudev T, Lim C. Effect of carboxylate-binding mode on metal binding/selectivity and function in proteins. Acc Chem Res. 2007 Jan;40(1):85–93. doi:10.1021/ar068181i PubMed PMID: 17226948.

23. Aurora R, Rose GD. Helix capping. Protein Sci. 1998 Jan;7(1):21–38. doi:10.1002/pro.5560070103 PubMed PMID: 9514257; PubMed Central PMCID: PMC2143812.

24. Dissmeyer N, Schnittger A. Use of phospho-site substitutions to analyze the biological relevance of phosphorylation events in regulatory networks. Methods Mol Biol. 2011;779:93–138. doi:10.1007/978-1-61779-264-9_6 PubMed PMID: 21837563.

25. Bilbrough T, Piemontese E, Seitz O. Dissecting the role of protein phosphorylation: a chemical biology toolbox. Chem Soc Rev. 2022 Jul 4;51(13):5691–730. doi:10.1039/d1cs00991e PubMed PMID: 35726784.

26. Balasuriya N, Kunkel MT, Liu X, Biggar KK, Li SSC, Newton AC, et al. Genetic code expansion and live cell imaging reveal that Thr-308 phosphorylation is irreplaceable and sufficient for Akt1 activity. J Biol Chem. 2018 Jul 6;293(27):10744–56. doi:10.1074/jbc.RA118.002357 PubMed PMID: 29773654; PubMed Central PMCID: PMC6036199.

27. Parks DH, Chuvochina M, Chaumeil PA, Rinke C, Mussig AJ, Hugenholtz P. A complete domain-to-species taxonomy for Bacteria and Archaea. Nat Biotechnol. 2020 Sep;38(9):1079–86. doi:10.1038/s41587-020-0501-8 PubMed PMID: 32341564.

28. Fakih F, Nandipati S, Kachale A, Heller J, Žváček J, Brázdovič F, et al. Frequent occurrence and predicted functions of tRNAs with 4-base-pair anticodon stems in bacteria: extended superwobble hypothesis. Nucleic Acids Res. 2026 Apr 24;54(7):gkag327. doi:10.1093/nar/gkag327.

29. Melnykov AV. New genetic codes in bacteria and archaea identified with a fast *k*-mer based algorithm. bioRxiv [Preprint]. 2026 [cited 2026 Apr 14]. p. 2026.04.02.715157. Available from: https://www.biorxiv.org/content/10.64898/2026.04.02.715157v1. doi:10.64898/2026.04.02.715157.

30. Grettenberger CL, Hamilton TL. Metagenome-assembled genomes of novel taxa from an acid mine drainage environment. Appl Environ Microbiol. 2021 Aug 11; 87(17):e00772–21. doi:10.1128/AEM.00772-21 PubMed PMID: 34161177; PubMed Central PMCID: PMC8357290.

31. Roda-Garcia JJ, Haro-Moreno JM, López-Pérez M. Evolutionary pathways for deep-sea adaptation in marine planktonic Actinobacteriota. Front Microbiol. 2023 May 10;14:1159270. doi:10.3389/fmicb.2023.1159270 PubMed PMID: 37234526.

32. Shulgina Y, Eddy SR. Codetta: predicting the genetic code from nucleotide sequence. Bioinformatics. 2022 Dec 13;39(1):btac802. doi:10.1093/bioinformatics/btac802 PubMed PMID: 36511586; PubMed Central PMCID: PMC9825746.

33. Simão FA, Waterhouse RM, Ioannidis P, Kriventseva EV, Zdobnov EM. BUSCO: assessing genome assembly and annotation completeness with single-copy orthologs. Bioinformatics. 2015 Oct 1;31(19):3210–2. doi:10.1093/bioinformatics/btv351 PubMed PMID: 26059717.

34. Lee RS, Pagan J, Wilke-Mounts S, Senior AE. Characterization of *Escherichia coli* ATP synthase beta-subunit mutations using a chromosomal deletion strain. Biochemistry. 1991 Jul 16;30(28):6842–7. doi:10.1021/bi00242a006 PubMed PMID: 1829962.

35. Burke B, Yang F, Chen F, Stehlin C, Chan B, Musier-Forsyth K. Evolutionary coadaptation of the motif 2−acceptor stem interaction in the class II prolyl-tRNA synthetase system. Biochemistry. 2000 Dec 1;39(50):15540–7. doi:10.1021/bi001835p.

36. Cusack S, Härtlein M, Leberman R. Sequence, structural and evolutionary relationships between class 2 aminoacyl-tRNA synthetases. Nucleic Acids Res. 1991 Jul 11;19(13):3489–98. doi:10.1093/nar/19.13.3489 PubMed PMID: 1852601; PubMed Central PMCID: PMC328370.

37. Aravind L, Koonin EV. DNA polymerase β-like nucleotidyltransferase superfamily: Identification of three new families, classification and evolutionary history. Nucleic Acids Res. 1999 Apr 1;27(7):1609–18. doi:10.1093/nar/27.7.1609.

38. Hasegawa T, Miyano M, Himeno H, Sano Y, Kimura K, Shimizu M. Identity determinants of *E. coli* threonine tRNA. Biochem Biophys Res Commun. 1992 Apr 15;184(1):478–84. doi:10.1016/0006-291x(92)91219-g PubMed PMID: 1567450.

39. Becker HD, Giegé R, Kern D. Identity of prokaryotic and eukaryotic tRNA^Asp^ for aminoacylation by aspartyl-tRNA synthetase from *Thermus thermophilus*. Biochemistry. 1996 Jan 1;35(23):7447–58. doi:10.1021/bi9601058.

40. Charron C, Roy H, Blaise M, Giegé R, Kern D. Non-discriminating and discriminating aspartyl-tRNA synthetases differ in the anticodon-binding domain. EMBO J. 2003 Apr 1;22(7):1632–43. doi:10.1093/emboj/cdg148 PubMed PMID: 12660169; PubMed Central PMCID: PMC152893.

41. Bailly M, Blaise M, Roy H, Deniziak M, Lorber B, Birck C, et al. tRNA-dependent asparagine formation in prokaryotes: characterization, isolation and structural and functional analysis of a ribonucleoprotein particle generating Asn-tRNA(Asn). Methods. 2008 Feb;44(2):146–63. doi:10.1016/j.ymeth.2007.11.012 PubMed PMID: 18241796.

42. Feng L, Yuan J, Toogood H, Tumbula-Hansen D, Söll D. Aspartyl-tRNA synthetase requires a conserved proline in the anticodon-binding loop for tRNA(Asn) recognition *in vivo*. J Biol Chem. 2005 May 27;280(21):20638–41. doi:10.1074/jbc.M500874200 PubMed PMID: 15781458.

43. Yamao F, Muto A, Kawauchi Y, Iwami M, Iwagami S, Azumi Y, et al. UGA is read as tryptophan in *Mycoplasma capricolum*. Proc Natl Acad Sci U S A. 1985 Apr;82(8):2306–9. doi:10.1073/pnas.82.8.2306 PubMed PMID: 3887399; PubMed Central PMCID: PMC397546.

44. McCutcheon JP, Moran NA. Functional convergence in reduced genomes of bacterial symbionts spanning 200 My of evolution. Genome Biol Evol. 2010;2:708–18. doi:10.1093/gbe/evq055 PubMed PMID: 20829280; PubMed Central PMCID: PMC2953269.

45. Parks DH, Chaumeil PA, Chuvochina M, Hugenholtz P. First report of stop codon reassignment to tryptophan in members of the bacterial phylum Actinomycetota. bioRxiv [Preprint]. 2025 [cited 2025 Dec 29]. p. 2025.12.01.691617. Available from: https://www.biorxiv.org/content/10.64898/2025.12.01.691617v1. doi:10.64898/2025.12.01.691617.

46. Hara KY, Kato-Yamada Y, Kikuchi Y, Hisabori T, Yoshida M. The role of the beta DELSEED motif of F1-ATPase: propagation of the inhibitory effect of the ε subunit. J Biol Chem. 2001 Jun 29;276(26):23969–73. doi:10.1074/jbc.M009303200.

47. Grosjean H, de Crécy-Lagard V, Marck C. Deciphering synonymous codons in the three domains of life: co-evolution with specific tRNA modification enzymes. FEBS Lett. 2010 Jan 21;584(2):252–64. doi:10.1016/j.febslet.2009.11.052 PubMed PMID: 19931533.

48. Ohama T, Inagaki Y, Bessho Y, Osawa S. Evolving genetic code. Proc Jpn Acad Ser B Phys Biol Sci. 2008 Feb;84(2):58–74. doi:10.2183/pjab.84.58 PubMed PMID: 18941287; PubMed Central PMCID: PMC2805505.

49. Moukadiri I, Garzón MJ, Björk GR, Armengod ME. The output of the tRNA modification pathways controlled by the *Escherichia coli* MnmEG and MnmC enzymes depends on the growth conditions and the tRNA species. Nucleic Acids Res. 2014 Feb;42(4):2602–23. doi:10.1093/nar/gkt1228 PubMed PMID: 24293650; PubMed Central PMCID: PMC3936742.

50. Emms DM, Kelly S. OrthoFinder: phylogenetic orthology inference for comparative genomics. Genome Biol. 2019 Nov 14;20(1):238. doi:10.1186/s13059-019-1832-y PubMed PMID: 31727128; PubMed Central PMCID: PMC6857279.

51. Nguyen LT, Schmidt HA, von Haeseler A, Minh BQ. IQ-TREE: a fast and effective stochastic algorithm for estimating maximum-likelihood phylogenies. Mol Biol Evol. 2015 Jan;32(1):268–74. doi:10.1093/molbev/msu300 PubMed PMID: 25371430; PubMed Central PMCID: PMC4271533.

52. Katoh K, Standley DM. MAFFT multiple sequence alignment software version 7: improvements in performance and usability. Mol Biol Evol. 2013 Apr;30(4):772–80. doi:10.1093/molbev/mst010 PubMed PMID: 23329690; PubMed Central PMCID: PMC3603318.

53. Capella-Gutiérrez S, Silla-Martínez JM, Gabaldón T. trimAl: a tool for automated alignment trimming in large-scale phylogenetic analyses. Bioinformatics. 2009 Aug 1;25(15):1972–3. doi:10.1093/bioinformatics/btp348.

54. Letunic I, Bork P. Interactive Tree of Life (iTOL) v6: recent updates to the phylogenetic tree display and annotation tool. Nucleic Acids Res. 2024 Jul 5;52(W1):W78–82. doi:10.1093/nar/gkae268 PubMed PMID: 38613393; PubMed Central PMCID: PMC11223838.

55. Mistry J, Chuguransky S, Williams L, Qureshi M, Salazar GA, Sonnhammer ELL, et al. Pfam: The protein families database in 2021. Nucleic Acids Res. 2021 Jan 8;49(D1):D412–9. doi:10.1093/nar/gkaa913 PubMed PMID: 33125078; PubMed Central PMCID: PMC7779014.

56. Eddy SR. Accelerated profile HMM searches. PLoS Comput Biol. 2011 Oct;7(10):e1002195. doi:10.1371/journal.pcbi.1002195 PubMed PMID: 22039361; PubMed Central PMCID: PMC3197634.

57. Seemann T. Prokka: rapid prokaryotic genome annotation. Bioinformatics. 2014 Jul 15;30(14):2068–9. doi:10.1093/bioinformatics/btu153 PubMed PMID: 24642063.

58. Rice P, Longden I, Bleasby A. EMBOSS: the European Molecular Biology Open Software Suite. Trends Genet. 2000 Jun;16(6):276–7. doi:10.1016/s0168-9525(00)02024-2 PubMed PMID: 10827456.

59. Mann HB, Whitney DR. On a test of whether one of two random variables is stochastically larger than the other. Ann Math Stat. 1947 Mar;18(1):50–60. doi:10.1214/aoms/1177730491.

60. Will S, Joshi T, Hofacker IL, Stadler PF, Backofen R. LocARNA-P: accurate boundary prediction and improved detection of structural RNAs. RNA. 2012 May;18(5):900–14. doi:10.1261/rna.029041.111 PubMed PMID: 22450757; PubMed Central PMCID: PMC3334699.

61. Ronquist F, Huelsenbeck JP. MrBayes 3: Bayesian phylogenetic inference under mixed models. Bioinformatics. 2003 Aug 12;19(12):1572–4. doi:10.1093/bioinformatics/btg180 PubMed PMID: 12912839.

62. Sullivan MJ, Petty NK, Beatson SA. Easyfig: a genome comparison visualizer. Bioinformatics. 2011 Apr 1;27(7):1009–10. doi:10.1093/bioinformatics/btr039 PubMed PMID: 21278367; PubMed Central PMCID: PMC3065679.

63. Camacho C, Coulouris G, Avagyan V, Ma N, Papadopoulos J, Bealer K, et al. BLAST+: architecture and applications. BMC Bioinformatics. 2009; 10:421. doi:10.1186/1471-2105-10-421 PubMed PMID: 20003500; PubMed Central PMCID: PMC2803857.

64. Tegenfeldt F, Kuznetsov D, Manni M, Berkeley M, Zdobnov EM, Kriventseva EV. OrthoDB and BUSCO update: annotation of orthologs with wider sampling of genomes. Nucleic Acids Res. 2025 Jan 6;53(D1):D516–22. doi:10.1093/nar/gkae987.

65. Sprinzl M, Horn C, Brown M, Ioudovitch A, Steinberg S. Compilation of tRNA sequences and sequences of tRNA genes. Nucleic Acids Res. 1998 Jan 1;26(1):148–53. doi:10.1093/nar/26.1.148.

